# Defects in the neuroendocrine axis cause global development delay in a *Drosophila* model of NGLY1 Deficiency

**DOI:** 10.1101/241653

**Authors:** Tamy Portillo Rodriguez, Joshua D. Mast, Tom Hartl, Ethan O. Perlstein

**Affiliations:** Perlara PBC, 6000 Shoreline Court, Suite 204, South San Francisco, California 94080, United States

## Abstract

N-glycanase 1 (NGLY1) Deficiency is a rare monogenic multi-system disorder first described in 2014. NGLY1 is evolutionarily conserved in model organisms, including the *Drosophila melanogaster* NGLY1 homolog, *Pngl*. Here we conducted a natural history study and chemical-modifier screen on a new fly model of NGLY1 Deficiency engineered with a nonsense mutation in *Pngl* at codon 420, resulting in truncation of the C-terminal carbohydrate-binding PAW domain. Homozygous mutant animals exhibit global development delay, pupal lethality and small body size as adults. We developed a 96-well-plate, image-based, quantitative assay of *Drosophila* larval size for use in a screen of the 2,650-member Microsource Spectrum compound library of FDA approved drugs, bioactive tool compounds, and natural products. We found that the cholesterol-derived ecdysteroid molting hormone 20-hydroxyecdysone (20E) rescued the global developmental delay in mutant homozygotes. Targeted expression of a human NGLY1 transgene to tissues involved in ecdysteroidogenesis, e.g., prothoracic gland, also rescues global developmental delay in mutant homozygotes. Finally, the proteasome inhibitor bortezomib is a potent enhancer of global developmental delay in our fly model, evidence of a defective proteasome “bounce-back” response that is also observed in nematode and cellular models of NGLY1 Deficiency. Together, these results demonstrate the therapeutic relevance of a new fly model of NGLY1 Deficiency for drug discovery, biomarker discovery, pharmacodynamics studies, and gene modifier screens.

## INTRODUCTION

Recessive loss-of-function mutations in the evolutionarily conserved gene NGLY1 result in an ultra-rare genetic disease called NGLY1 Deficiency, which is characterized by global developmental delay, seizures, small head and extremities, chronic constipation, lack of tears, and floppy body (Enns et al., 2014). NGLY1, short for N-glycanase 1, encodes a deglycosylating enzyme that hydrolyzes N-linked glycans from asparagine residues of glycoproteins, liberating oligosaccharides for degradation and recycling (Suzuki et al., 2016). A comprehensive clinical snapshot by National Institutes of Health (NIH) established potential measurable clinical endpoints and a baseline of disease progression in a cohort of 12 patients (Lam et al., 2017).

The pathophysiology of NGLY1 Deficiency has not yet been fully resolved. Before 2014, little was known about NGLY1 function. Its elucidation is the focus of numerous research groups employing a diversity of disease models, including model organisms and human cells. Two models of the pathogenesis of NGLY1 Deficiency have been proposed.

The first model is rooted in biochemistry and defects in cellular glycoprotein homeostasis (Huang et al., 2015). NGLY1 is an essential component of the conserved protein quality control system known as endoplasmic-reticulum-associated degradation (ERAD), bridging p97/VCP-mediated retrotranslocation of proteins from the ER to the cytoplasm for bulk deglycosylation and subsequent degradation by the ubiquitin-proteasome system (UPS) (Suzuki, 2015). The absence of cytoplasmic N-glycanase activity has been proposed to result in inappropriate cleavage of N-glycans from glycoproteins by the downstream cytoplasmic enzyme endo-β-*N*-acetylglucosaminidase (ENGase), whose normal substrate is soluble oligosaccharide liberated by NGLY1. Such glycoproteins misprocessed by ENGase would retain a single GlcNac residue that may destabilize proteins and promote their aggregation. Suzuki and colleagues observed evidence of *N*-GlcNac misprocessing and accumulation *in vitro* in NGLY1^-/-^ mouse embryonic fibroblasts (Huang et al., 2015; Fujihira et al., 2017). Based on the collective work of Suzuki and colleagues, inhibition of ENGase has been proposed as a therapeutic thesis for NGLY1 Deficiency (Fujihira et al., 2017; Bi et al., 2017). Indeed, a NGLY1^-/-^ ENGase^-/-^ double mutant mouse is viable while a NGLY1^-/-^ single mutant displays varying degrees of lethality depending on genetic background (Fujihira et al., 2017). However, a NGLY1^-/-^ ENGase^-/-^ double mutant mouse is not healthy. Additional pathogenic mechanisms are required to explain NGLY1 Deficiency.

The second model of pathogenesis is rooted in genetics and defects in the deglycosylation of specific glycoprotein clients, including but not limited to the master regulator of the conserved 26S proteasome “bounce-back” response, NFE2L1 (the transcription factor nuclear factor erythroid 2 like 1 also known as Nrf1) (Radhakrishnan, 2010). Nrf1, belongs to the ancient basic leucine zipper family of transcription factors that regulate many developmental and stress response pathways in animals (Kim, 2016). The fly Nrf1 homolog, *cap-n-collar (cnc)* increases the expression of proteasome subunit genes, as well as oxidative and redox stress response pathways (Grimberg, 2011). In an unbiased screen for genetic modifiers of the proteasome bounce-back response in nematodes, Lehrbach and Ruvkun discovered that the nematode homolog of NGLY1, PNG-1, deglycosylates the ER-membrane bound isoform of the nematode homolog of Nrf1, SKN-1A. They further demonstrated that deglycosylation of SKN-1A by PNG-1 is required for SKN- 1A translocation to the nucleus, and transcriptional activity (Lehrbach, 2016).

In a complementary study, Bertozzi and colleagues revealed that NGLY1 activity is required for Nrf1 signaling in mouse embryonic fibroblasts mice in the same manner that PNG-1 activity is required for SKN-1A function in nematodes (Tomlin, 2017). In fact, they showed that chemical inhibition of NGLY1 function potentiates cytotoxicity caused by proteasome inhibition in human cancer cell lines (Tomlin, 2017), which mirrors the observation in nematodes that png-1 loss-of-function mutants are extremely hypersensitive to proteasome inhibition by bortezomib (Lehrbach, 2016). Jafar-Nejad and colleagues showed that the fly NGLY1/Pngl is required during embryonic and larval development in *Drosophila* for post-translational processing of *Dpp*, the fly homolog of the conserved bone morphogen protein (BMP) family (Galeone, 2017), opening up the possibility that NGLY1 is required for the function of multiple glycoprotein clients.

Here we carried out phenotyping, high-throughput assay development and a chemical-modifier screen on a new fly model of NGLY1 Deficiency, herein referred to as ngly1^PL^. Of ~2,650 bioactive compounds, the ecdysteroid insect molting hormone 20-hydroxyecdysone (20E) suppressed global developmental delay in mutant homozygotes. Expression of a human NGLY1 transgene in the prothoracic gland (PG) and sites of ecdysteroidogenesis rescued global developmental delay in mutant homozygotes. These data indicate that cell autonomous defects in ecdysone-producing organs and tissues may explain the global developmental delay of the ngly1^PL^ flies. ngly^PL^ flies are also hypersensitive to the proteasome inhibitor bortezomib and the organic solvent dimethyl sulfoxide (DMSO). Together, these observations can be parsimoniously accommodated by a model wherein NGLY1*/Pngl* is required for Nrf1/cnc function in the *Drosophila* neuroendocrine axis.

## METHODS

### ngly1^PL^ allele generation

A cassette containing a stop codon and *mw+* was cloned into a modified version of pUC57. Homology arms to the *ngly1* locus were cloned upstream and downstream of the cassette. The guide RNA (GCTGAGGAATAACTTTCGAT CGG) was cloned into pCDF3 (Port et al., 2014). pCDF3 and pUC57 with *ngly1* homology arms, stop codon, and *mw+* were injected into *vas-Cas9* (Bloomington stock #51323). Two independent *mw+* F1 strains were established and balanced stocks were created. Sequencing confirmed the integration of the stop codon and *mw+* (Figure 1A) immediately 3' to bp 1906699 (Release 5.57) in chromosome 2R (NT_033778.3).

**Figure 1.**
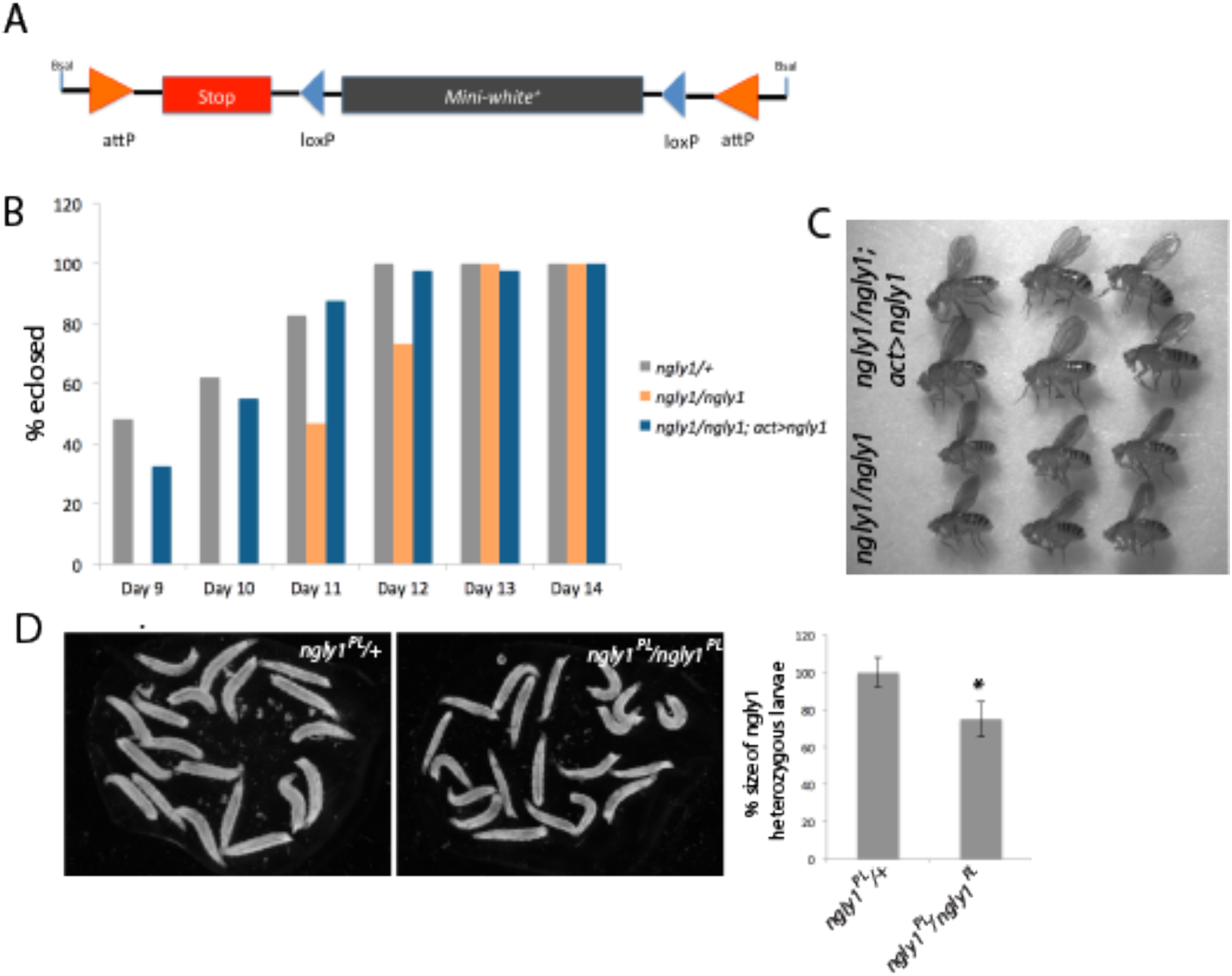
Genotyping and phenotyping *ngly1^PL^* allele. **A)** We used CRISPR to create an allele of ngly1 with a stop codon and white + transgene inserted upstream of the PAW domain. Sequencing confirmed the integration of the stop codon and *mw+* immediately 3’ to bp 1906699 (Release 5.57) in chromosome 2R (NT_033778.3). **B)** Ubiquitous expression of a human *ngly1* transgene rescued the 2-day development delay to eclosion phenotype observed on *ngly1^PL^* homozygotes. **C)** *ngly1^PL^* homozygous adult flies are smaller than their homozygous siblings expressing the human *ngly1* transgene. **D)** *ngly1^PL^* homozygous are developmentally delayed, at 3 days old, the larvae are 25% smaller than their heterozygous siblings (p<0.01).

### Rate of eclosion and rescue by human ngly1 transgene studies

*ngly1PL/CyO; actin-switch-gal4; UAS-human-ngly1, w+/+* males were crossed to ngly1PL/CyO, Act-GFP virgin females. The phenotypes of eclosing flies were recorded on days 9-14 after parents were mated. We found that the actin-switch-gal4 transgenic strain expressed Gal4, despite the absence of RU486.

### Fly strains

Actin5C-switch-gal4 (stock #9431) was obtained from the Bloomington stock center. Pngl excision alleles and *UAS-human-ngly1* were obtained from Dr. Hamed Jafar-Nejad. 2_286, phm and spok GAL4 drivers were obtained from Dr. Michael O'Connor.

### Larval size measurements

0-4 hour old larvae were placed into petri dishes with standard fly food media for 3 days at 25°C. Then, larvae were removed from the food, rinsed in PBS, and placed in PBS to be imaged. The areas of nineteen heterozygous and twenty homozygous larvae were measured in ImageJ.

### Plate preparation for chemical modifier screening

100nL compound or DMSO (vehicle) was dispensed from mother plates into wells of 96 well daughter plates using the Echo acoustic dispenser (LabCyte). Then, 100uL of molten standard fly food media (molasses, agar, yeast, propionic acid) lacking cornmeal, but carrying 0.025% bromophenol blue, was dispensed using a MultiFlo (Bio-Tek Instruments). Bromophenol blue dye in the guts of the larva increases their signal over background later during imaging. Plates were placed on a plate shaker and shaken for 1 minute, during which, the molten agar solidified with thoroughly mixed compound/DMSO in each well. Plates were then covered with adhesive aluminum seals and stored at 4°C for up to two weeks.

### Culturing fly larvae in 96-well plates

*Ngly1PL/CyO, Act-GFP* flies were cultured in a large population cage (Genessee) for up to two weeks, where they laid eggs on grape juice agar trays (Genessee) coated with a thin strip of active yeast paste. Egg collections were restricted to ~6 hours during morning hours and were placed into 25°C for ~24 hrs. ~0-6hr old 1^st^ instar larvae were rinsed off of the trays with room temperature water and collected in funnel-attached 40 micron sieves typically used for cell straining. To remove embryo contamination, the larvae/embryo mixture was incubated two times in 10.3% inoculation solution (sodium chloride, sucrose, 10% Triton X-100) for three minutes. Most embryos float to the top of this solution after three minutes, while larvae remain in suspension. Embryos from the mixture's solution can therefore be removed with a 10ml pipette. Embryos were then added to sorting solution (Polyethylene glycol, COPAS 200x GP sheath reagent, Tween20) and drawn into a BioSorter (Union Biometrica) for sorting and dispensing into 96 well compound/media bearing plates. Three GFP negative homozygotes were gated from heterozygotes and dispensed into the 11 left most columns of the plate, and wells G12 and H12. Three GFP positive heterozygotes were dispensed into wells A12, B12, C12, D12, E12, and F12. Plates were then covered with permeable adhesive seals and incubated at 25°C for three days. Approximately 18 plates were dispensed into per day, and larval viability was not affected by their time submerged in dispensing solution (<4 hours).

### Preparation of plates for imaging

On the terminal day of the assay (day 3), gas permeable seals were removed and plates were filled with 20% sucrose solution made acidic with hydrochloric acid (pH 2) and carrying defoamer (Five Star Defoamer 105-2 oz). Plates were then vortexed and more solution was added to suspend larvae to a fixed focal plane. Plates were then imaged./

### Image and data analysis

The Fly Imager uses a Sony a7r ii camera, controlled over USB by the gphoto software to generate full plate images that are then run through a larval detection algorithm. The algorithm first builds an image of what the well would look like when it is empty. To do this it excludes areas near edges (since those are probably larvae) and interpolates across the gaps. It then looks at the difference between the image and estimated empty background, selecting areas with a high difference to be larvae.

### 20Efeeding and effect on developmental timing

20E (Enzo) was dissolved in 100% ethanol and added to molten standard fly food media at 200μM and added to vials. An equivalent set of vials with an equal amount of ethanol was added for negative controls. 0-6 hour larvae were dispensed with the BioSorter into the vials and incubated at 25°C for 12 days. The number of larvae that pupariated were recorded on key days at 8am, 12pm, and 4pm, and the number of adults that eclosed were recorded on days 9 – 12 at a single time.

### Rescue of developmental delay with ring and prothoracic gland expression of human ngly1

Ring gland expression of human *ngly1* was driven using the GAL4/UAS system (Brand and Perrison, 1993) and ring gland drivers 286-GAL4, spok-GAL4, and phm-GAL4 in 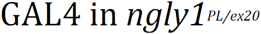 compound heterozygotes. Adults were allowed to lay eggs for 8-16 hours, and the genotypes for all eclosing flies were recorded. The number of ring gland rescued flies eclosed was normalized to the number of eclosed sibling homozygotes not carrying the driver.

## RESULTS

### Generation of the ngly1^PL^ fly and characterization of its phenotypes

We used CRISPR/Cas9 to create a new allele of *Pngl* with a premature stop codon and a selectable marker inserted upstream of the PAW domain, herein referred to simply as “ngly1^PL^” (**Fig. 1A**). We benchmarked ngly1^PL^ against the previously described *Pngl* genetic null mutant generated by P-element excision *(Pngl^ex20^)*, which causes developmental defects and semi-lethality with few adult escapers (Funakoshi, 2010). ngly1^PL^ homozygotes are delayed in the larval-to-pupal transition by one day, and delayed to eclosion by 2 days (**Fig. 1B**). As three-day-old larvae, ngly1^PL^ homozygotes are ~75% the size of their heterozygote siblings (*P* < *1×10^-10^, student's t-test)* (**Fig. 1D**). Consistent with the semi-lethality of *Pngl* excision alleles, we also observed that the ngly1^PL^ homozygote mutant exhibits cold-sensitive pupal lethality, meaning no survival below 18°C (data not shown). Time to eclosion in ngly1^PL^ homozygotes is completely rescued by ubiquitous expression of a human NGLY1 (hNGLY1) transgene (**Fig. 1B**). The small body size phenotype of ngly1^PL^ homozygote adults is also rescued by ubiquitous expression of hNGLY1 (**Fig. 1C**). These results when flies are reared in standard vials suggested phenotypes suitable for high-throughput, whole-organism phenotypic screening at each stage of fly development from 1^st^ instar larvae onwards.

We decided to optimize a high-throughput larval size assay in 96-well plates for several reasons. Assay miniaturization from 30mL vials to 96-well plates involves reducing the number of animals tested by a factor of 10, e.g., 30 animals per vial versus 3 animals per well. The larval size difference between ngly1^PL^ heterozygote larvae versus ngly1^PL^ homozygote larvae was more statistically robust in 96-well plates than the timing to pupation difference or the timing to eclosion difference. A 3-day assay versus a 7-11 day assay allowed for faster optimization cycle times. Time to pupation and time to eclosion would be secondary assays in vial format to ensure primary screening hits rescue global developmental delay, not just developmental delay in larvae.

As a prelude to drug screening, we established the tolerability of ngly1^PL^ homozygote larvae to dimethyl sulfoxide (DMSO), the organic solvent for almost every compound library, including the Microsource Spectrum collection. The maximum tolerated dose of DMSO, therefore, sets a ceiling on the final assay screening concentration. Unexpectedly, we observed that ngly1^PL^ homozygotes are extremely hypersensitive to DMSO compared to the heterozygote control (**Figure 2**). We estimated a maximum tolerated dose in the ngly1^PL^ homozygote of 0.1% v/v DMSO (14mM) (**Fig. 2A, C**), which is actually several fold more sensitive than *Pngl* excision allele 20 (**Fig. 2B**). In comparison, wild-type *Drosophila* exhibit adult lethality starting at 1% v/v DMSO, larval lethality ranging from 2% v/v to 3% v/v DMSO, and a “no observed adverse effect level” (NOAEL) dose of 0.3% v/v DMSO (Nazir, 2003). DMSO hypersensitivity appears to be specific to the ngly1^PL^ homozygote in a limited comparison to other mutants (data not shown). For the purposes of drug screening, we exploited DMSO as a sensitizer in the larval size assay. Exposure of mutant larvae to 0.1% v/v DMSO, which would entail a final assay screening concentration of 10μM for each library compound, was potent enough to increase the dynamic range and improve the robustness of the assay while sparing larval lethality.

**Figure 2.**
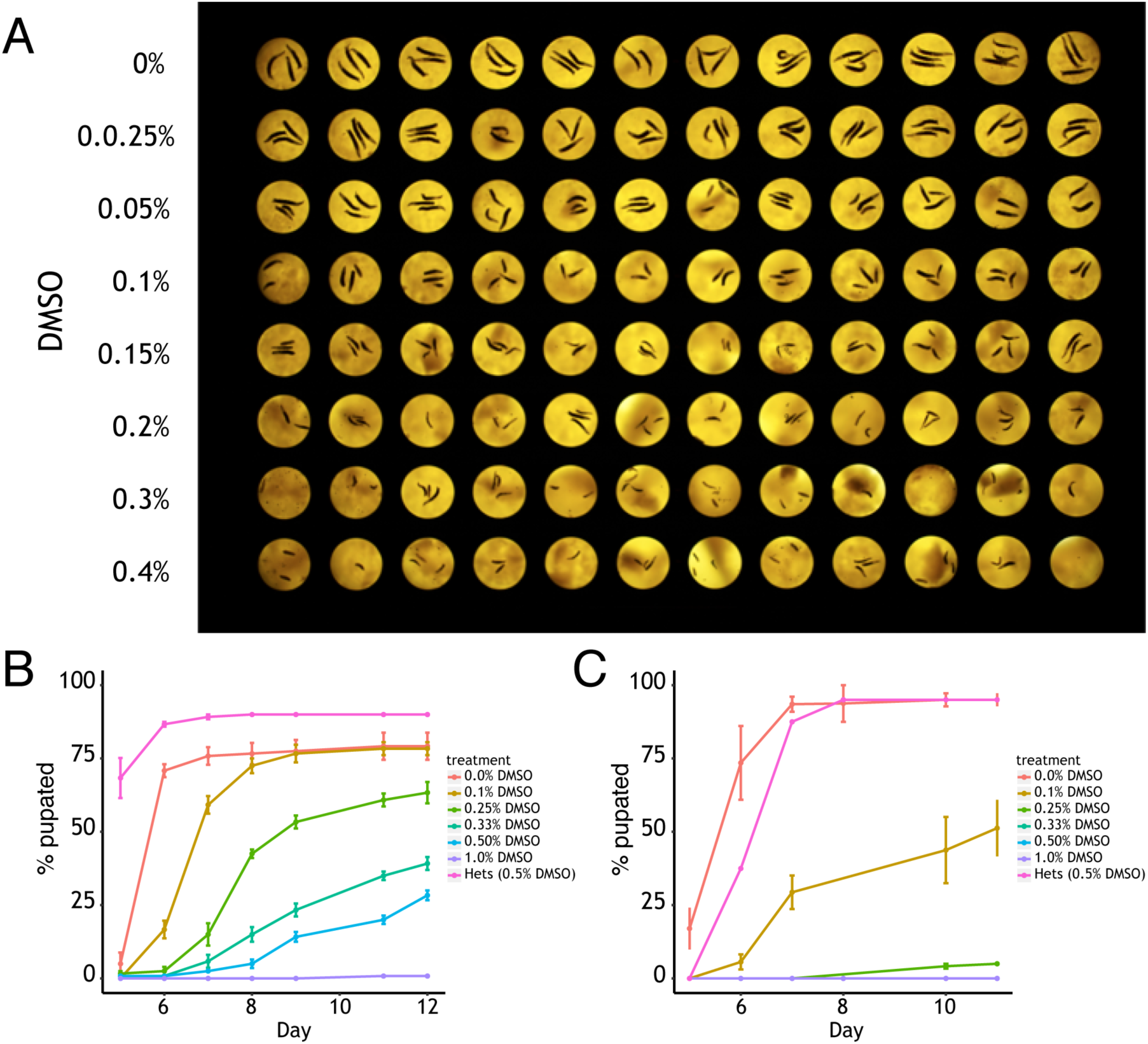
ngly1^PL^ homozygotes are hypersensitive to dimethyl sulfoxide (DMS**O)**. **A)** An image of *ngly1^PL^* homozygous larvae grown in 96-well plate format on food dosed with different concentrations (0%-0.4%) of DMSO. *ngly1^PL^* homozygous larvae are hypersensitive to DMSO, a decrease in larval size is observed starting at 0.025% DMSO and larval size is decrease as the dose of DMSO is increased. The time to pupation of **B)** *ngly1^Ex20^* or **C)** *ngly1^PL^* homozygous larvae reared on food treated with DMSO. Homozygous mutant larvae exhibit hypersensitivity to DMSO showing a delayed time to pupariate and increased lethality.

### A statistically robust high-throughput image-based larval size chemical modifier pilot screen yielded one validated hit, 20-hydroxyecdysone (20E)

We posited that the small larval size phenotype of ngly1^PL^ homozygote mutants could be used to discover small-molecule suppressors that provide insight into the pathophysiology of NGLY1/*Pngl* deficiency in the fly. To that end, we developed a novel image-based assay to culture *Drosophila* larvae in clear-bottom 96-well plates where each well either contains a unique small molecule or vehicle. Using a 96-well-plate format allowed us to take advantage of existing lab automation for managing multi-well plates in high-throughput screening (HTS) applications, including a whole-organism sorter and dispenser. Our method includes steps to dissolve and dilute pre-existing fly food so that larvae can be floated to a fixed focal plane and then imaged with custom plate imager. Software was written to measure the overall area of floated larvae to enable the identification of small molecules that significantly increase the size of ngly1^PL^ homozygote larvae vs vehicle-fed larvae.

We selected the Microsource Spectrum collection for a pilot screen. The library contains 2,532 unique compounds including ~600 FDA approved drugs, ~800 compounds that have reached clinical trial stages in the USA, ~400 drugs that have been marketed in Europe or Japan but not the USA, ~600 bioactive tool compounds, and ~800 natural products. Three larvae per well were cultured in 96-well plates with control wells comprising the two outermost columns. We used ngly1^PL^ homozygous larvae fed vehicle (0.1% v/v DMSO) as a negative control. We include two positive control groups: the first group consist of ngly1^PL^ heterozygous larvae fed vehicle, and the second group consists of ngly1^PL^ homozygous larvae cultured without DMSO. The remaining 80 wells contained ngly1^PL^ homozygous larvae fed a unique library compound at 10^M plus 0.1% v/v DMSO. An image of a representative drug screening plate with zoomed-in reference wells is shown in **Figure 3**.

**Figure 3.**
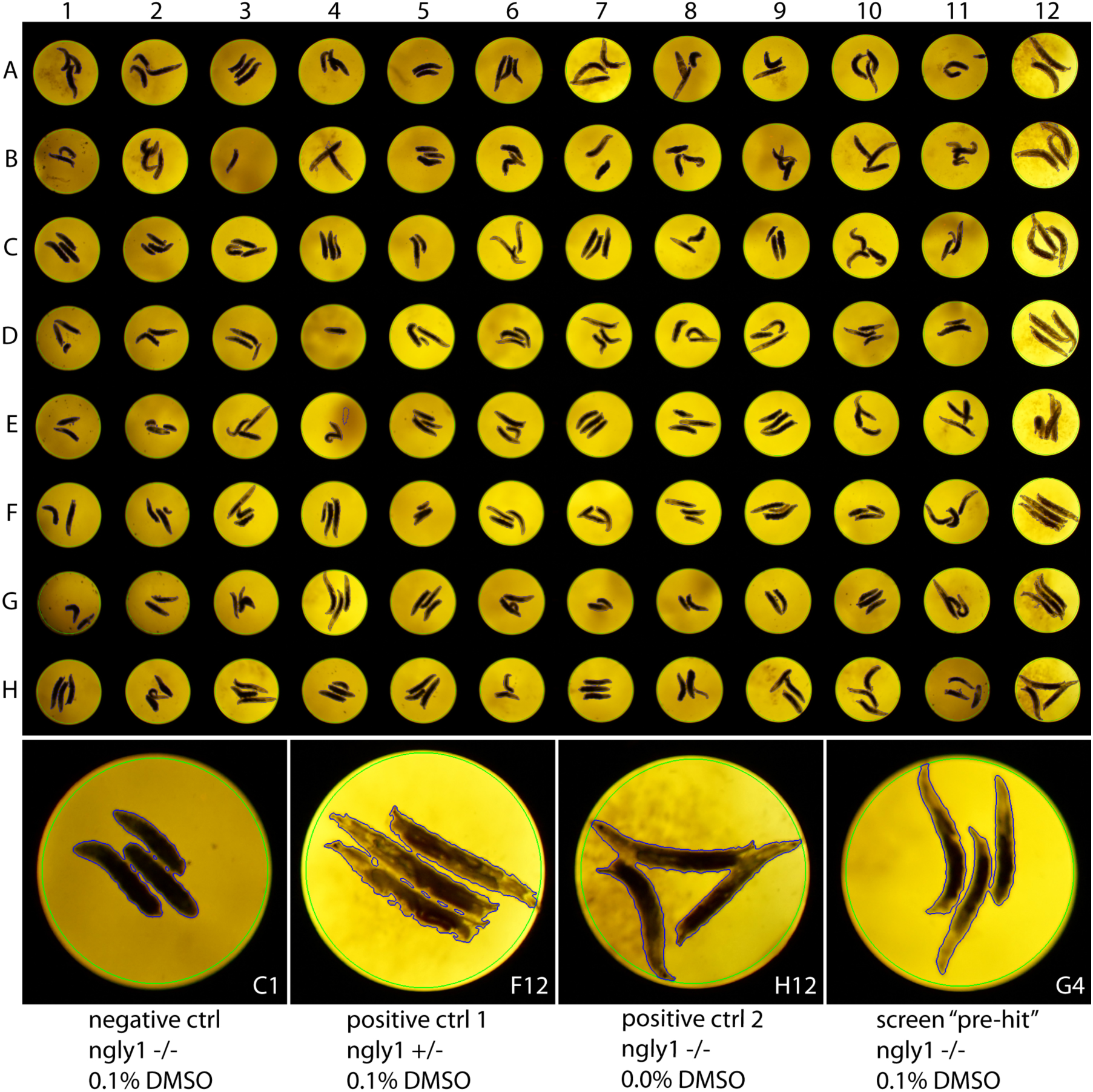
Layout of an *nglyl* assay plate with examples of controls and pre-hit compound. Larvae were cultured in 96-well plates with negative controls on the left most wells (column 1) which carried *ngly1^PL^* homozygous with vehicle DMSO at 0.1%. 80 testing wells were in the middle (columns 2–11) which carried *ngly1^PL^* homozygous larvae with a compound at 10uM, and DMSO at 0.1%. Right most top wells (12A-12**F)** contain positive control group 1 with *ngly1^PL^* heterozygous larvae and 0.1% DMSO. Right most bottom wells (12G-12**H)** contain positive control group 2 with *ngly1^PL^* homozygous larvae without DMSO. The compound in well G4 was considered a pre-hit compound.

We performed the screen in triplicate, meaning three independent biological replicates. There was a robust statistical separation between positive and negative controls (**Fig. 4A**). On average, ngly1^PL^ homozygotes were half the size of ngly1^PL^ heterozygotes, although some variation in size was observed in each genotype. 75% of all screening plates (73 of 96) had a Z' factor > 0; most of the screening plates with high variability belonged to the second replicate (**Fig. 4B**). A weak positive correlation existed in pairwise comparisons of each biological replicate when all data points are included (**Fig. 5A**). When we only included wells with Z scores less than −2 or greater than 2, the positive correlation increased significantly on average (**Fig. 5B**). For example, in the pairwise comparison of replicate 1 versus replicate 2, the correlation improves from R^2^ = 0.05977 in the full dataset to R^2^ = 0.48171 in the reduced dataset (with positive and negative controls removed as well).

**Figure 4.**
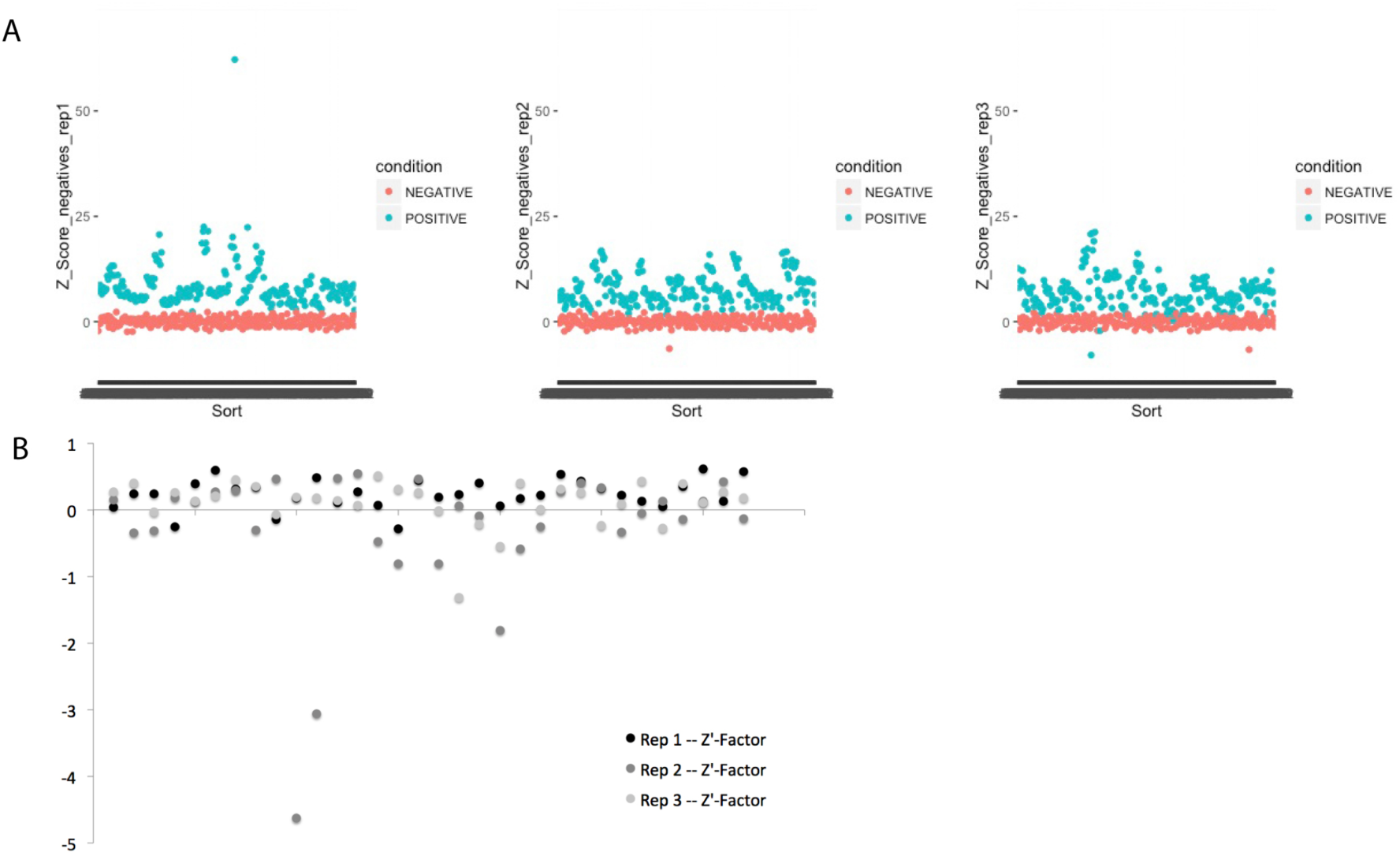
The assay had a consistent difference between positive and negative controls. **A)** A consistent separation between positive and negative controls with Z'Factor >0, indicated a robust assay that could identify chemical suppressors. **B)** Three replicates of a 32 plate library were analyzed and 73 of 96 plates had a Z'Factor >0 which further suggest a robust assay to identify chemical suppressors.

**Figure 5.**
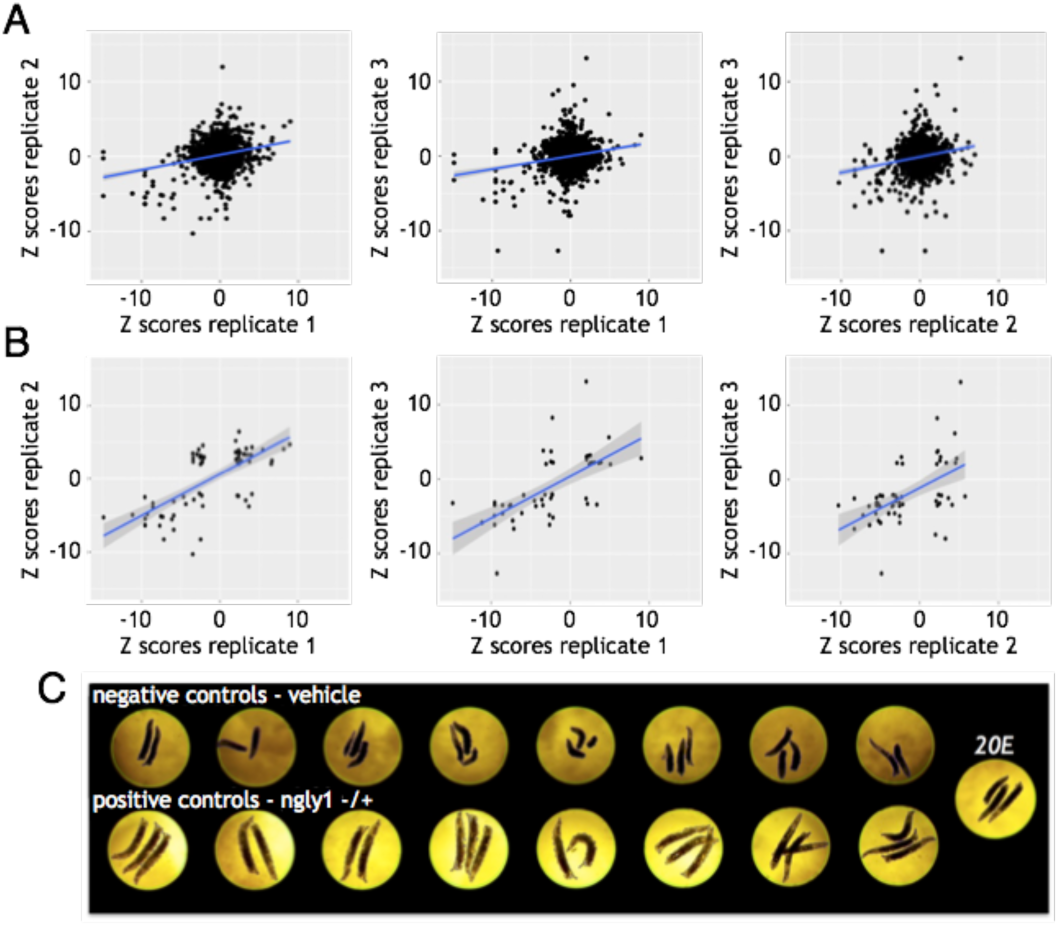
Positive correlation between 3X replicates. **A)** The three pairwise comparisons of Z-scores show positive correlations indicating that a set of small molecules could modify the small larval size phenotype **B)** The positive correlations between replicates is more evident when only plotting Z-scores of < −2 or > 2. **C)** Larval size is rescued when 20E is fed to *ngly1^PL^* homozygous larvae.

We initially considered 162 pre-hits with a Z score of > 2 in two of three biological replicates as primary screening positives (Table 1). Over two-thirds of those pre-hits proved to be false positives for one of two of the following reasons. First, uneaten fly food in the well occasionally increased the image background artificially inflating the calculated area of the larvae. Second, because *Pngl* mutants are hypersensitive to DMSO, any failure in compound dispensing or variability in DMSO levels due to hydration resulted in larger larvae. Forty-five compounds with a Z-score of > 2 in two of three replicates were considered further because their wells did not have obvious high background or low/no DMSO. The raw images of the wells containing those 45 compounds were manually inspected, and 22 had a clear difference from controls. Ultimately, 22 compounds were found to have a Z of > 2 in two of three replicates and upon manual inspection, the wells with those compounds appeared to have larvae larger than the negative controls in the same plate. Two of those 22 compounds had a Z of > 2 in three of three replicates, were counted twice in this comparison, and thus the final set of pre-hits we tested further consists of 18 unique compounds.

**Table 1.** Whittling screen pre-hits to a set of promising compounds to test in validation studies. Ultimately, 22 compounds were found to have a Z of > 2 in 2 of 3 replicates and upon manual inspection, the wells with those cpds appeared to have larvae larger then the negative controls. Two of those 22 compounds had a Z of > 2 in 3 of 3 replicates, were counted twice in this comparison, and thus the final set to consider further were 18 unique compounds.

|  | Compounds with Z > 2<br>in 2 of 3 replicates | High background | Low/No DMSO | Without high background<br>or Low/No DMSO | Clear difference from controls<br>found by manual inspection |
| --- | --- | --- | --- | --- | --- |
| *Replicates 1 and 3* | 44 | 30 | 3 | 11 | 5 |
| *Replicates 1 and 2* | 42 | 12 | 5 | 25 | 12 |
| *Replicates 2 and 3* | 76 | 64 | 3 | 9 | 5 |
| *Total* | 162 | 106 | 11 | 45 | 22 |

**Table 2.**
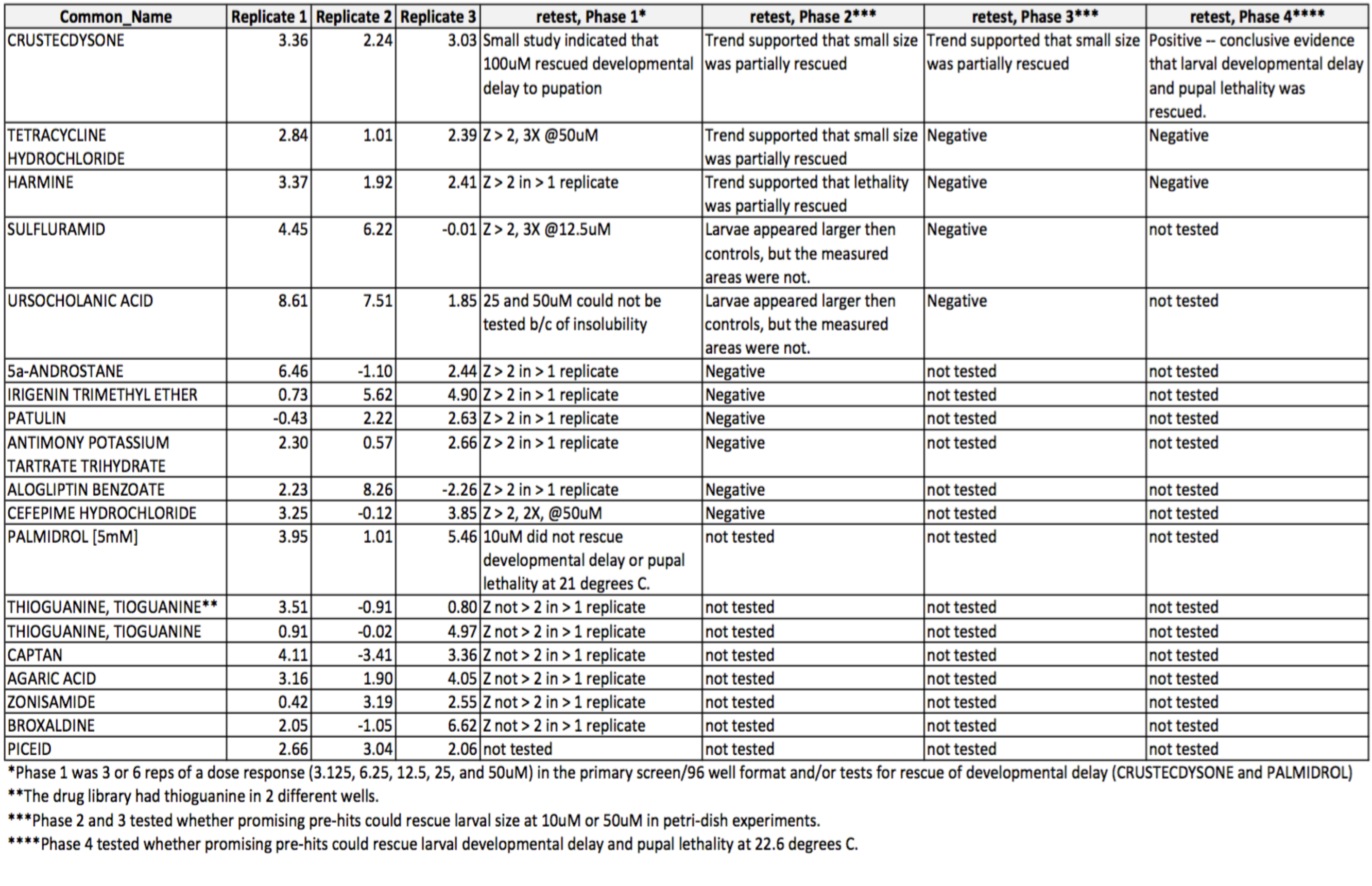
The 18 compounds that showed promise in the screen and were considered pre-hits. These had a Z-score of >2 in 2 of 3 replicates in the primary screen.

All 18 of the pre-hits from the screen were ordered, dissolved, and retested in attempt to reproduce rescue in vial format with larger numbers of animals (**Fig. 5C**, Table 2). One compound, 20-hydroxyecdysone (20E), rescued the developmental delay of ngly1^PL^ homozygote larvae development to pupae (**Fig. 6B**), but had no effect on developmental timing of ngly1^PL^ heterozygote larvae (**Fig. 6A**). Moreover, the suppressive effect of 20E persisted to adulthood, resulting in a statistically significant 4-fold increase in eclosion percentage (**Fig. 6E**). As a control to rule out a simple ecdysone biosynthesis defect, we showed that the 20E precursor 7-hydroxycholesterol (7D) failed to rescue ngly1^PL^ homozygote larvae development to pupae (**Fig. 6D**). If synthesis of 20E is faulty, it is likely at a point downstream of 7D. We could not reproducibly validate any of the other 17 pre-hits, so we focused our efforts on understanding the mechanism-of-action (MoA) of 20E. Therefore, this pilot screen had an extremely low hit rate of 0.04% (1/2532).

**Figure 6.**
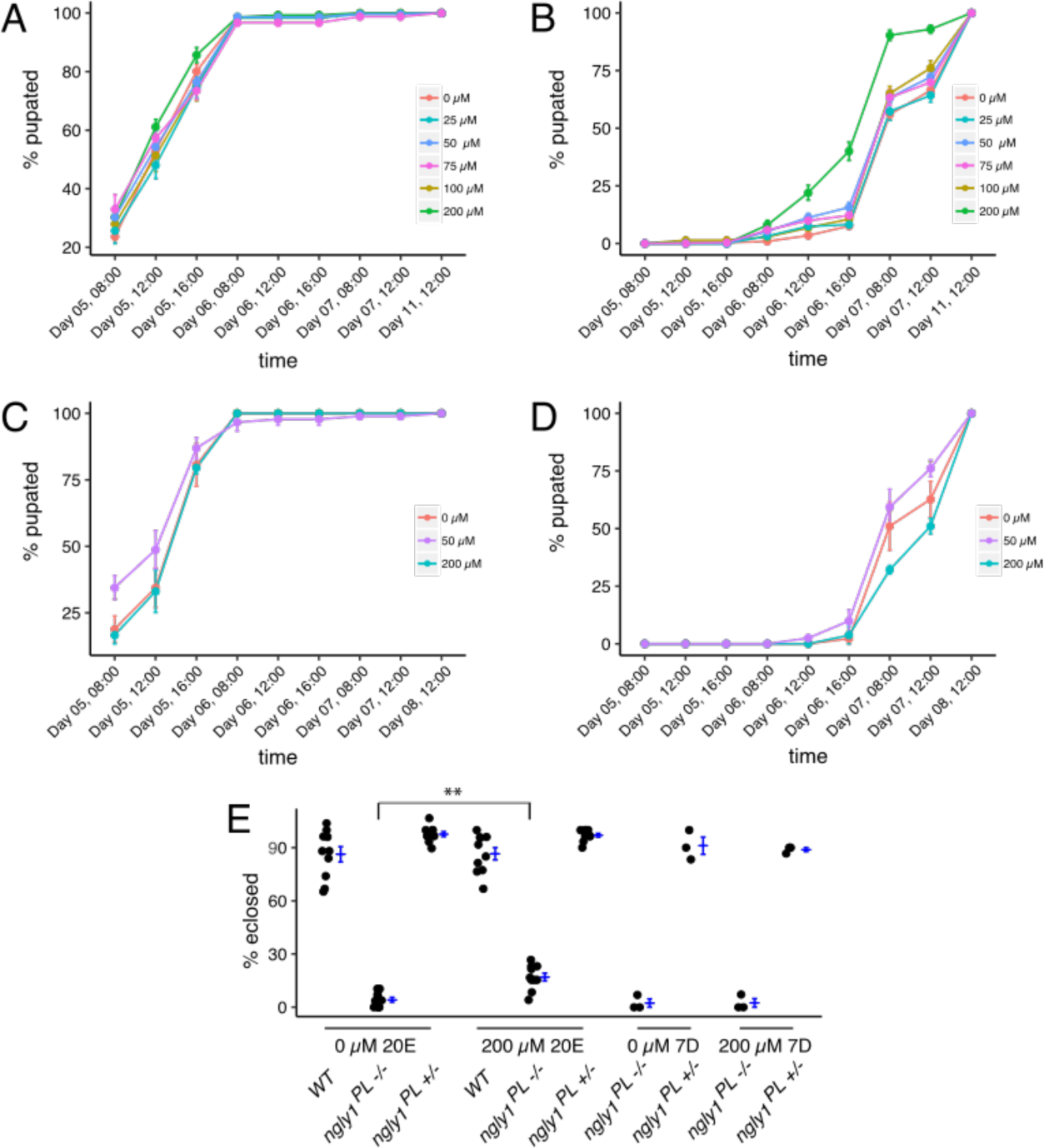
20-hydroxyecdysone (20E), but not an earlier ecdysone pathway precursor (7-dehydrocholesterol), partially rescues developmental delay and lethality in *ngly1^PL^ mutants*. The time to pupation of **A)** *ngly1^PL^* heterozygous or **B)** *ngly1^PL^* homozygous larvae reared on food treated with different concentration of 20E. 20E did not impact developmental rate of heterozygous *ngly1^PL^* larvae to pupation, but rescued development delay of homozygous *ngly1^PL^* larvae to pupation at 200μM. The time to pupation of **C)** *ngly1^PL^* heterozygous or **D)** *ngly1^PL^* homozygous larvae reared on food treated with different concentration of 7-d. 7-d did not impact developmental rate of heterozygous or homozygous *ngly1^PL^* larvae to pupation. **E)** The fraction of animals surviving to eclosion. 20E, but not 7-d, rescued larval lethality of *ngly1^PL^* homozygous at 200uM.

### 20E implicates the fly neuroendocrine axis as particularly sensitive to loss of NGLY1/Pngl function

20E drives metamorphosis in *Drosophila* and arthropods generally (Faunes, 2016). Dietary cholesterol forms the basis of 20E, and all 20E precursors are synthesized in the prothoracic gland, an organ of < 50 cells that comprises part of the larger ring gland. The immediate 20E precursor, ecdysone or “E”, is packaged into secretory vesicles, secreted, and distributed by the hemolymph throughout the animals. E is converted to 20E in these peripheral tissues, and initiates signaling cascades and gene expression inducing physiological, morphological, and behavioral changes with each molt, or developmental transition.

To test whether the 20E insufficiency in Pngl-deficient mutants is autonomous to the ring gland, we expressed a UAS-driven hNGLY1 transgene that can rescue global developmental delay when expressed ubiquitously (**Fig. 1B, C**), in the ring gland with the 2-286-GAL4 driver. To control for off-target mutations due to strain background confounding our results, we tested the trans-heterozygous combination of *Pngl* alleles, ngly1^PL^/ex20. Most ngly1^PL^/ex20 larvae not expressing the human ngly1 transgene eclosed on Day 10. In contrast, most ngly1^PL^/ex20 larvae expressing human NGLY1 in the ring gland pupated on Day 9 (**Fig. 7A**). Aside from the ring gland, 2-286-GAL4 also drives expression in the salivary gland, fat body, and cuticle in larvae (Timmons et al., 1997). We observed a similar rescue effect when human *ngly1* transgene expression is driven by two ring-gland-specific drivers, *phantom-GAL4 (phm)* (**Fig. 7B**) and *spookier-GAL4 (spok)* (**Fig. 7C**). *phm* and *spok* encode cytochrome P450 monooxygenases in the ecdysone biosynthetic pathway (Rewitz, 2007). Together, these data indicate that NGLY1/*Pngl* is necessary for normal function of the ring gland to enable proper levels of 20E to circulate throughout the developing animal and drive developmental transitions.

**Figure 7:**
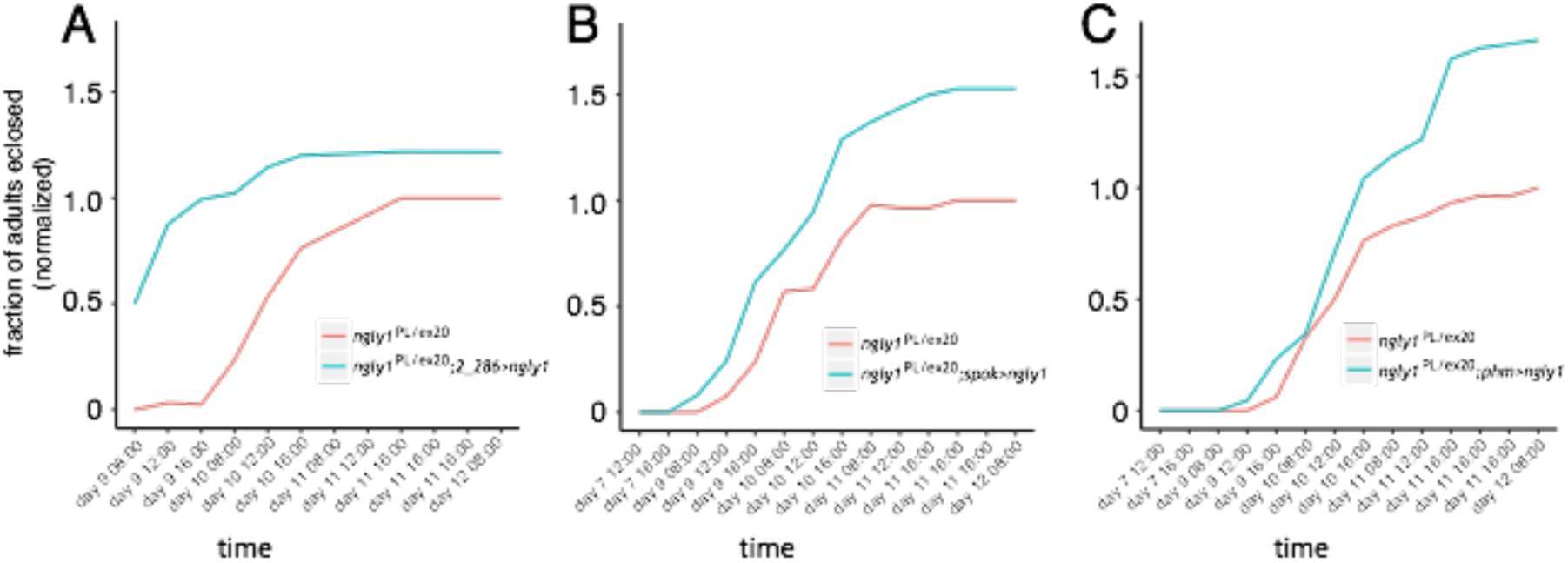
*nglyl* is necessary for normal function of the ring gland. The fraction of *ngly1^PL^ / ngly1^Ex20^* compound heterozygotes with human *nglyl* driven by the ring gland driver (blu**e) A)** *2_286-GAL4* **B)** *spokier-GAL4*, or **C)** *phantam-GAL4* surviving to eclosion compared to sibling controls lacking a driver (red). **A)** Compound heterozygous *ngly1^PL^ / ngly1^Ex20^* larvae expressing *2_286>ngly1* eclosed earlier and had lower lethality than control flies. Compound heterozygous *ngly1^PL^ / ngly1^Ex20^* larvae expressing *spokier>ngly1* or *phantam>ngly1* eclosed lower lethality than control flies.

### NGLY1/Pngl deficient flies are hypersensitive proteasome inhibition

As mentioned above, loss of NGLY1 sensitizes nematodes and human cancer cell lines to proteasome inhibition. We predicted that ngly1^PL^ homozygote mutants would exhibit hypersensitivity to bortezomib. Indeed, bortezomib caused 100% lethality of ngly1^PL^ homozygous larvae at 5μM, while lethality was not observed in ngly1^PL^ heterozygous larvae until a dose of 25μM (**Figure 8**.). In addition, the size of ngly1^PL^ homozygous mutants was reduced by <50% when treated with 1μM bortezomib, and to ~50% ngly1^PL^ heterozygous larvae at ~10μM bortezomib.

**Figure 8.**
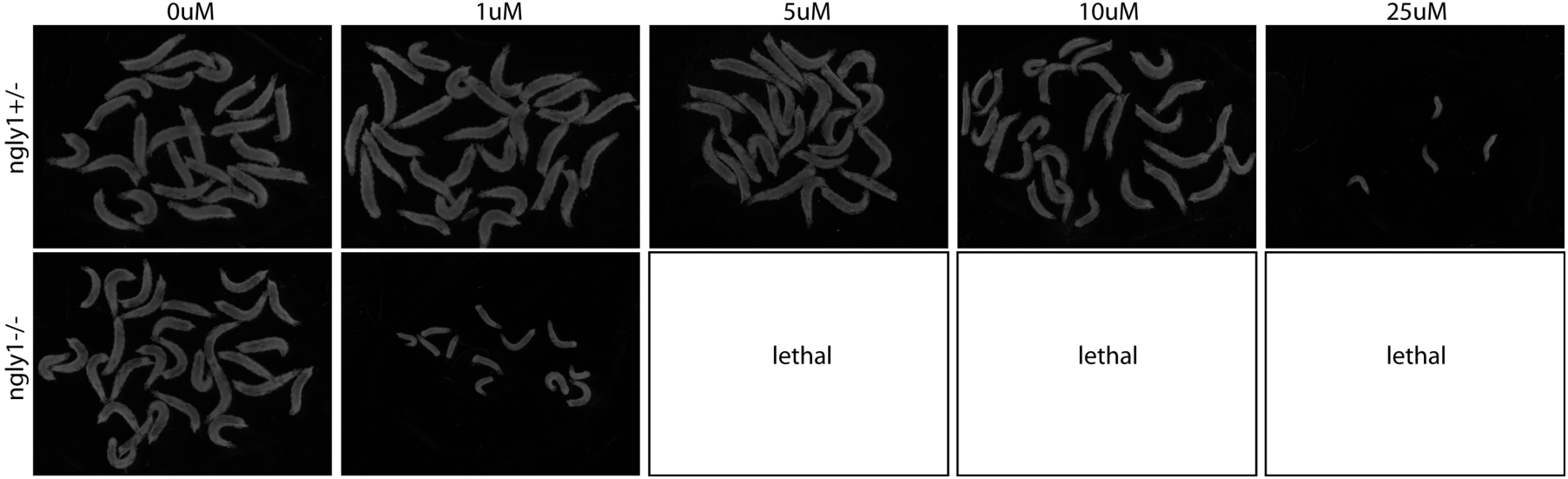
NGLY1 larvae are ≥10X more sensitive to the proteasome inhibitor bortezomib. Images of 3 day *ngly1^PL^+^/-^* heterozygous or age matched *ngly1^PL-/-^* mutant larvae raised on food treated with 0, 1, 5, 10 or 25μM bortezomib. Bortezomib delays larval developmental progression in *ngly1^PL^* homozygous larvae more severely than *ngly1^PL^* heterozygous larvae leading to smaller sized larvae, and is lethal for homozygotes at concentrations equal to or greater than 5 μM.

These data indicate that the half-maximal inhibitory concentration (IC50) of bortezomib to reduce larval growth is ~10X less in ngly1^PL^ homozygote.

## DISCUSSION

We successfully generated a new *Drosophila* model of NGLY1 Deficiency, optimized a high-throughput whole-animal phenotypic assay of larval size, and then demonstrated therapeutic relevance in a proof-of-concept drug repurposing screen. While 20E itself should not be considered a drug candidate for NGLY1 Deficiency in humans, the fact that it is a chemical suppressor implicates the neuroendocrine axis in the pathophysiology of *Pngl* deficiency in flies. In other words, even though 20E is an insect-specific developmental hormone, the neuroendocrine axis and steroid-derived developmental hormones are conserved in mammals and may play a role in NGLY1 Deficiency in humans. Collectively, our findings - most strikingly, hypersensitivity of the ngly1^PL^ mutant both to bortezomib and to DMSO - align with results observed in nematodes (Lehrback and Ruvkun, 2017) and human cells (Tomlin, 2017) that point to the essential and conserved role of NGLY1 in regulating the function of glycoprotein clients, specifically Nrf1 and the proteasome bounce-back response.

In fact, we can already propose a mechanism to link Nrf1 function to ecdysteroidogenesis and the neuroendocrine axis in flies. The fly homolog of Nrf1 is the longest isoform of *cnc, CncC*, which contains a conserved N-terminal leucine rich transmembrane region targeting *CncC* to the ER (Grimberg, 2011). Specific loss of *CncC* in the prothoracic gland reduces the expression of ecdysone biosynthetic genes and results in delayed timing to pupation (Deng and Kerppola, 2013). RNA interference of the Colorado potato beetle homolog of *CncC* also reduced ecdysteroidogenesis pathway gene expression and delayed timing pupation, which could be rescued by 20E supplementation (Sun et al., 2017). Our data make a strong testable prediction: *CncC* is the functional equivalent of SKN-1A in nematodes and Nrf1 in mammals, and therefore is a substrate for deglycosylation by *Pngl* in flies. A second testable prediction is that the fly homolog of nematode DDI-1 and human DDI2, *rings lost (ringo)*, acts downstream of *Pngl* to proteolyze *CncC*, generating a mature, nuclear-active species.

There are other explanations for how 20E might rescue global developmental delay in the ngly1^PL^ mutant that do not directly involve Nrf1/CncC, or that may act in concert with loss of Nrf1/CncC activity. First, ngly1^PL^ mutants may not package ecdysone into secretory vesicles properly, or may be defective in secreting ecdysone. Second, signal transduction mediated by the interaction between 20E and the ecdysone receptor (EcR) might not be fully operational, and so extra 20E boosts this flawed signaling. Third, ecdysone secretion by the ring gland is induced by two neurons that synapse onto the prothoracic gland and secrete the neuropeptide prothoracicotropic hormone, or PTTH. PTTH contacts the receptor tyrosine kinase *Torso* to initiate signaling leading to ecdysone secretion. Perhaps there is a flaw in PTTH secretion or *Torso* signaling. Fourth, damage to developing larval tissues, or starvation, may impinge on 20E to fine tune organism development so that a properly proportioned and nourished animal can develop fully to adulthood. The ngly1^PL^ mutant may have damaged tissues, for example through protein-aggregate toxicity or proteasome stress; or it may have some degree of starvation, for example if the gut cannot attain nutrients properly. 20E feeding may bypass delays induced by this hypothetical tissue damage/starvation.

## ACKNOWLEDGMENTS

We thank the Grace Science Foundation for funding this work. We thank Dr. Kevin Lee for planning and experiment planning. Drs. Hamed Jafar-Nejad and Tadashi Suzuki kindly provided reagents. Peter Sand designed and built our custom fly imager and analysis software. Dr. Tom Lee provided assistance during ngly1^PL^ mutant strain construction. Dr. Rebecca Choy provided early guidance on how to scale up fly husbandry/culturing.

